# From Synchrony to Asynchrony: Cerebellar-Basal Ganglia Functional Circuits in Young and Older Adults

**DOI:** 10.1101/309344

**Authors:** Hanna K. Hausman, T. Bryan Jackson, James R.M. Goen, Jessica A. Bernard

**Author notes:** Corresponding Author: Jessica A. Bernard, 515 Coke Street, 4235 TAMU, College Station, TX 77843-4235. Denotes shared first-authorship. These authors contributed equally to this work.

## Abstract

Resting state functional magnetic resonance imaging (rs-fMRI) has indicated disruptions in functional connectivity in older (OA) relative to young adults (YA). While age differences in cortical networks are well studied, differences in subcortical networks are poorly understood. Both the cerebellum and the basal ganglia are of particular interest given their role in cognitive and motor functions, and work in non-human primates has demonstrated direct reciprocal connections between these regions. Here, our goal was twofold. First, we were interested in delineating connectivity patterns between distinct regions of the cerebellum and basal ganglia, known to have topologically distinct connectivity patterns with cortex. Our second goal was to quantify age-differences in these cerebellar-striatal circuits. We performed a targeted rs-fMRI analysis of the cerebellum and basal ganglia in 33 YA and 31 OA individuals. In the YA, we found significant connectivity both within and between the cerebellum and basal ganglia, in patterns supporting semi-discrete circuits that may differentially subserve motor and cognitive performance. We found a shift in connectivity, from one of synchrony in YA, to asynchrony in OA, resulting in substantial age differences. Connectivity was also associated with behavior. These findings significantly advance our understanding of cerebellar-basal ganglia interactions in the human brain.

## Introduction

Even in healthy aging, older adults (OA) experience declines in motor and cognitive functions (e.g., Salthouse 1996; Reuter-Lorenz et al. 1999; Park et al. 2001; Woollacott and Shumway-cook 2002; Heuninckx et al. 2005; Seidler et al. 2010; Anguera et al. 2011). Concurrent with these performance declines are changes in both brain structure (e.g., Raz et al. 1998, 2005, 2013; Salat et al. 2005; Walhovd et al. 2011; Bernard and Seidler 2013) and functional activation patterns (Reuter-Lorenz et al. 1999; Cabeza 2002; Davis et al. 2008; Seidler et al. 2010). While functional MRI (fMRI) has provided critical insights into the aging mind and brain, the fMRI testing environment itself may negatively affect performance due to scanner noises and participant stress responses to a novel environment (Hommel et al. 2012; Koten et al. 2013; van Maanen et al. 2016). This may exacerbate age effects in performance. Resting state fMRI (rs-fMRI) provides an alternative approach and allows for the examination of fluctuations in brain activity at rest (e.g., Biswal et al. 1995, 2010). In the absence of a task, brain regions that typically activate together during task performance show strong correlations with one another. Importantly, this methodology has delineated subcortical connectivity networks of the cerebellum (Habas et al. 2009; Krienen and Buckner 2009; O’Reilly et al. 2010; Bernard et al. 2012, 2014) and basal ganglia (Di Martino et al. 2008).

rs-fMRI is particularly useful for the study of aging and clinical populations. As noted above, this methodology removes the confounds of task performance in the scanner, and the scan times are on average, relatively short (from 5 to 15 minutes). The short scan times are beneficial for participant comfort, particularly in groups that may present with symptoms (physical or psychological) that may make the scanning environment less comfortable. Further in aging and clinical populations, the lack of task demands is particularly beneficial. In advanced age, cognitive and motor performance deficits are not uncommon, and having to complete a task in a novel environment, while lying on one’s back with images presented through mirror goggles may further confound performance. Investigating resting state networks removes this potential confound, while still providing important insight into functional brain organization. Across a now fairly extensive literature on rs-fMRI in OA, typically, there are decreases across the board in network connectivity in older adults relative to younger adults (e.g., Andrews-Hanna et al. 2007; Damoiseaux et al. 2008; Langan et al. 2010; Bernard et al. 2013; Bo et al. 2014). Notably, these age differences in functional connectivity are also related to behavioral performance (both motor and cognitive) in OA (e.g., Andrews-Hanna et al. 2007; Langan et al. 2010; Bernard et al. 2013), supporting the functional relevance of these differences. Most of the rs-fMRI studies to date have investigated the connectivity patterns of the default mode network due to its implications in Alzheimer’s disease (Ferreira and Busatto 2013). Subcortical network differences are relatively understudied, with few exceptions (Bernard et al. 2013; Bo et al. 2014).

The cerebellum is an important, but understudied, region in aging research. The cerebellum plays a role in both motor and cognitive behavior (e.g., Schmahmann and Sherman 1998; Desmond et al. 2005; Stoodley and Schmahmann 2009; E et al. 2012; Ferrucci et al. 2013). It is both smaller in volume (MacLullich et al. 2004; Raz et al. 2005a, 2013; Bernard and Seidler 2013; Miller et al. 2013) and has weaker functional connectivity in OA (Bernard et al., 2013), negatively affecting performance in motor and cognitive tasks (e.g., Bernard et al., 2013; Bernard & Seidler, 2013). Similarly, the basal ganglia contribute to diverse functional domains (reviewed in Bernard et al. 2017) and have connectivity patterns with the cortex that likely underlies these behavioral contributions (Di Martino et al. 2008; Draganski et al. 2008). Functional networks of the basal ganglia are also known to be different across the lifespan (Bo et al. 2014). Notably, however, there are also bi-directional direct connections between the cerebellum and the basal ganglia (Hoshi et al. 2005; Bostan and Strick 2010, 2018, Bostan et al. 2010, 2013). More recently, using tractography, connections between the cerebellum and basal ganglia have been mapped *in vivo* in the human brain as well (Pelzer et al. 2013; Milardi et al. 2016). These connections include direct disynaptic connections between the cerebellum and subthalamic nucleus (Bostan et al., 2010; Hoshi et al., 2005), but also connections through the thalamus (Milardi et al., 2016; Bostan & Strick, 2018). The direct connections and resulting communication between these regions are of great interest. Though the nature of the shared communication and functional role of these connections is only speculative at this point, there is a general consensus within the field that a systems-level approach to investigating these structures and interactions is of great importance, and that these interactions are of both behavioral and clinical significance (Caligiore et al. 2017).

Here, our goals were two-fold. First, we were interested in carefully characterizing patterns of rs-fMRI connectivity between the cerebellum and basal ganglia. Prior work from our own group and others has suggested that resting signal in the cerebellum and basal ganglia is significantly correlated (Di Martino et al. 2008; Habas et al. 2009; Bernard et al. 2012). However, to our knowledge, there has been no detailed investigation to date quantifying the connectivity patterns between lobules of the cerebellum and sub-regions of the basal ganglia. Here, we used the striatal seeds defined by DiMartino and colleagues (2008), along with spherical seeds placed in the cerebellar lobules, localized using the SUIT atlas (Diedrichsen 2006; Diedrichsen et al. 2009). We focused on Lobules V and VI, as well as Crus I and II, as these regions are associated with primary motor, motor preparatory, frontal, and parietal regions of the cortex (Kelly and Strick 2003; Krienen and Buckner 2009; Strick et al. 2009; Bernard et al. 2012) and have networks that are overlapping with those of the striatum (Di Martino et al. 2008). This approach allowed us to test the hypothesis that lobules of the cerebellum and subregions of the basal ganglia that are associated with similar cortical regions (Di Martino et al. 2008; Draganski et al. 2008; Habas et al. 2009; Krienen and Buckner 2009; Stoodley and Schmahmann 2009; Bernard et al. 2012; Bernard, Russell, et al. 2017; Stoodley et al. 2012) would show significant connectivity at rest. That is, regions of the cerebellum associated with motor networks and function would be correlated with regions of the basal ganglia that are thought to be similarly engaged.

Second, we sought to characterize age differences in these patterns of connectivity between the cerebellum and basal ganglia. Given that both structures contribute to the performance of motor and cognitive tasks (e.g., Imamizu et al. 2000; Doyon et al. 2002; Chen and Desmond 2005; Stoodley and Schmahmann 2009; Bernard et al. 2017), domains which show marked performance differences in older adults (e.g., Park et al. 2001; Seidler et al. 2010; Bernard et al. 2013), an understanding of differences in the interactions between these regions may provide important new insights into age-related performance declines. Indeed, our prior work has suggested that there is weaker connectivity between the cerebellum and basal ganglia, and that connectivity between these regions contributes to performance in older adults (Bernard et al. 2013). A detailed analysis of age differences in connectivity provides an important foundation for future work investigating these associations with respect to behavior and for investigations of disease impacting the basal ganglia and cerebellum, particularly Parkinson’s Disease (e.g., Wu and Hallett 2013; Bostan and Strick 2018). This also stands to provide important new insights into subcortical differences in OA, which are relatively understudied. We predicted that older adults would show lower connectivity between the regions and in connections that stand to impact both motor and cognitive behavior.

## Method

### Participants

We recruited 33 YA (age ± stdev; 22.47 ± 2.81 years, 14 females) and 31 OA (age ± stdev; 73.26 ± 5.25, 17 females) from the greater College Station community with fliers and emails. All participants are healthy individuals with no history of neurological disorders or stroke and had no contraindications for fMRI scanning. In each age group, two participants were left handed. This was due to the initial inclusion of left-handed OA. Two left-handed YA participants were then recruited for matching purposes. Notably, the proportion of left-handed individuals (6.66%) is just below what is typically seen in the population at large (approximately 10%) (Oldfield 1971), making our investigation more broadly generalizable. Prior to beginning all study procedures, participants signed a consent form approved by Texas A&M University Institutional Review Board. In addition, all participants completed the Montreal Cognitive Assessment (MOCA) (Nasreddine et al. 2005), as well as the Activities-Specific Balance Confidence (ABC) Scale (Powell and Myers 1995).

### rs-fMRI Acquisition

Rs-fMRI data were collected using a 3-tesla Siemens Verio scanner with a 12-channel head coil at the Texas A&M Institute for Preclinical Studies. A 7 minute and 18 second blood-oxygen-level dependent (BOLD) scan was acquired with an echo-planar functional protocol (number of volumes =165, repetition time [TR]=2620 ms, echo time [TE]= 29 ms; flip angle [FA]=75°, 3.8×3.8×3.5 mm3 voxels; 33 slices, field of view (FOV)= 240 x 240 mm). In addition, a high-resolution T1-weighted 3D magnetization prepared rapid gradient multi-echo sequences (MPRAGE; sagittal plane; TR=1900 ms; TE=2.99 ms; 0.9 mm3 isomorphic voxels; 160 slices; FOV=230 x 230 mm; FA=9°; time=5:59 minutes) to facilitate normalization of the rs-fMRI data.

### rs-fMRI Preprocessing and Analysis

Anatomical and structural images were pre-processed and analyzed using the Conn toolbox version 18b (Whitfield-Gabrieli and Nieto Castañón 2012). This included preprocessing procedures that were implemented in Conn using SPM. We followed a standard preprocessing pipeline which included functional realignment and unwarping, functional centering of the image to (0, 0, 0) coordinates, slice-timing correction, structural centering to (0, 0, 0,) coordinates, structural segmentation and normalization to MNI space, functional normalization to MNI space, and spatial smoothing with a smoothing kernel of 6mm FWHM, paralleling recent work from our group (Bernard, Goen, et al. 2017). Data were denoised using a band-pass filter of .008 - .09 Hz. White matter and cerebral-spinal fluid signals were linearly regressed out prior to first level analysis. Because of the potential confounding effects of motion and signal outliers (Power et al. 2012; Van Dijk et al. 2012) these procedures also included processing using the Artifact Rejection Toolbox (ART). This was set using the 97^th^ percentile settings and allowed for the quantification of participant motion in the scanner and the identification of outliers based on the mean signal. With these settings, the global-signal z-value threshold was set at 9mm, while the subject-motion threshold was set at 2 mm. 6-axis motion information and frame-wise outliers were included as covariates in our subsequent first-level analyses. The 6 motion covariates were reduced to one variable by averaging the absolute value of each axes’ average and the frame-wise time-series was averaged across each participant, resulting in one value for each measure for each participant for group comparison. Notably, the two groups differ on both measures, with OA showing increased movement overall (frame-wise outliers: YA: *M* = .18, *SD* = .06, OA: *M* = .27, *SD* = .11, *t*(61) = 2.68, *p* = .009; motion correction: YA: *M* = .09, *SD* = .04, OA: *M* = .14, *SD* = .09, *t*(61) = 4.36, *p* < .001). Data were then denoised using a band-pass filter of .008 - .09 Hz.

Rs-fMRI analysis focused on regions of interest (ROIs) in both the basal ganglia and cerebellum. The basal ganglia seed regions were based on those used by Di Martino and colleagues (2008) in their comprehensive mapping of striatal resting state connectivity, while cerebellar seeds were placed in the center of lobules V, VI, Crus I and Crus II as defined by the SUIT atlas (Diedrichsen, 2006; Diedrichsen et al., 2009). For all ROIs we created spherical seeds with a radius of 3.5mm. To limit multiple comparisons, we investigated cerebellar seeds in the right hemisphere and striatal seeds in the left hemisphere. Coordinates for the basal ganglia and the cerebellar seeds can be found in Table 1.

**Table 1:**
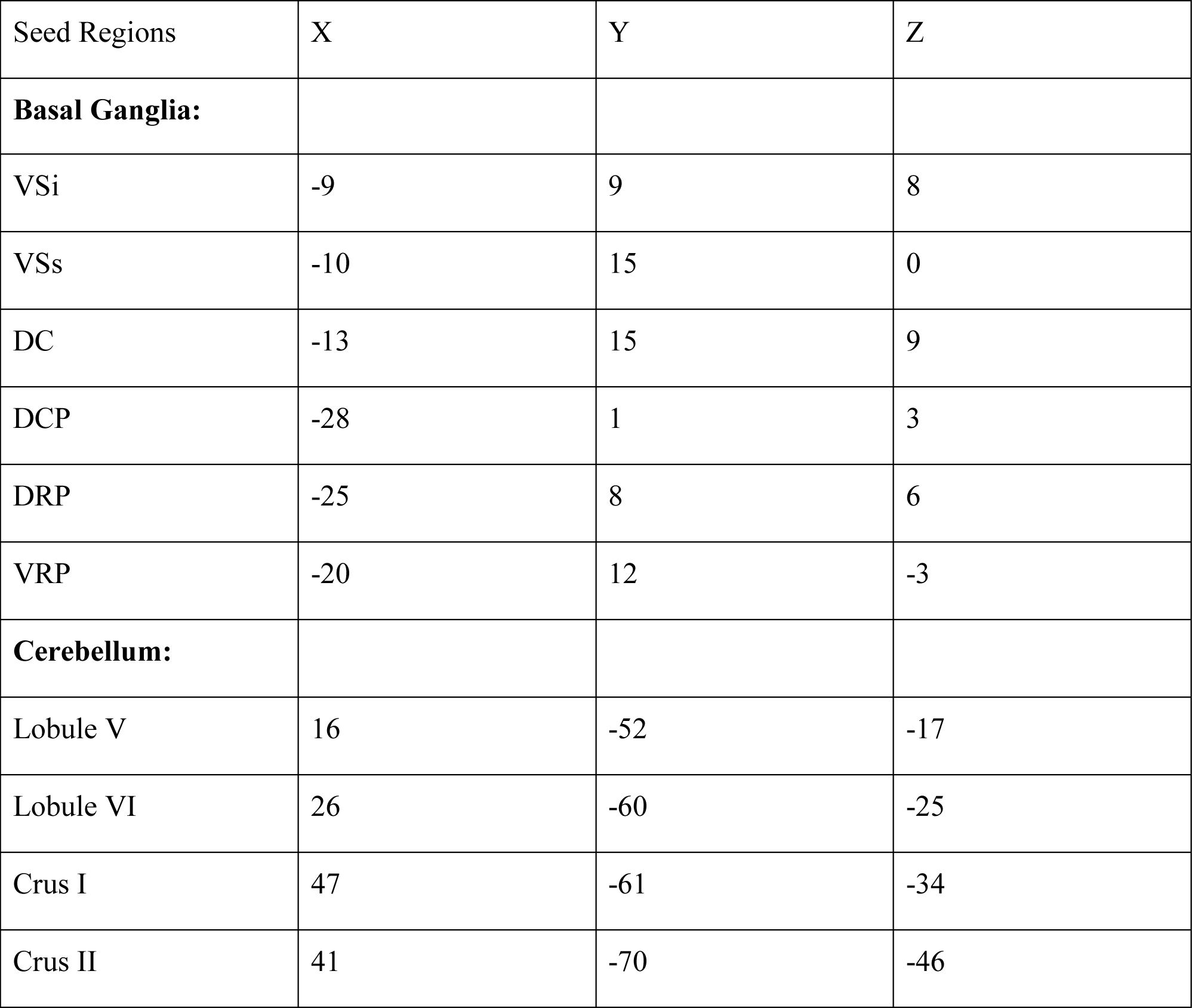
Coordinates for Striatal and Cerebellar Regions of Interest. VSi: inferior ventral striatum; VSs: superior ventral striatum; DC: dorsal caudate; DCP: dorsal caudal putamen; DRP: dorsal rostral putamen; VRP: ventral rostral putamen.

| Seed Regions | X | Y | Z |
| --- | --- | --- | --- |
| **Basal Ganglia:** |  |  |  |
| VSi | -9 | 9 | 8 |
| VSs | -10 | 15 | 0 |
| DC | -13 | 15 | 9 |
| DCP | -28 | 1 | 3 |
| DRP | -25 | 8 | 6 |
| VRP | -20 | 12 | -3 |
| **Cerebellum:** |  |  |  |
| Lobule V | 16 | -52 | -17 |
| Lobule VI | 26 | -60 | -25 |
| Crus I | 47 | -61 | -34 |
| Crus II | 41 | -70 | -46 |

We completed ROI-ROI bivariate correlation analysis to quantify the associations in activation at rest between subregions of the cerebellum and basal ganglia. We investigated within-group connectivity patterns of our seeds of interest, and we compared connectivity of the YA and OA groups in these regions. Correlation analyses were conducted directly in the Conn Toolbox using the Fisher transformed connectivity values (z scores). Results were evaluated using an analysis level FDR correction such that p<.05. This correction takes into account the total number of pairwise correlations run in our ROI-ROI analysis, and computes an FDR correction accordingly. This included a total of 90 comparisons. With 10 total seeds, the analysis computed all pairwise comparisons, with the exception of correlations of a seed to itself. While we are interested primarily in the correlations between the cerebellum and basal ganglia, the analysis also included correlations within these structures. Further, for all analyses we implemented permutation testing using 1000 permutations. We also used CONN to investigate correlations between ROI-ROI connectivity strength and behavioral performance to begin to quantify and understand the behavioral relevance of these networks. We looked at associations between connectivity and MOCA scores, a measure of cognitive ability, and with ABC scores, a self-report measure of balance confidence. Importantly, we assessed the relationships between these self-report measures and all of the ROI connections, including corrections for multiple comparisons. This allowed us to test the relative specificity of the connectivity-behavior associations.

Lastly, in addition to the analyses designed to quantify connectivity between cerebellar and basal ganglia ROIs, we were also interested in determining the relative functional segregation of these nodes and age differences therein. This allowed us to further interrogate the hypothesis that cerebellar and basal ganglia sub-regions that are part of similar cortical networks would be correlated with one another. Here we looked at the strength of connectivity between motor-motor pairings, cognitive-cognitive pairings, and cross-modal motor-cognitive pairings. Our “motor” nodes are Lobule V, Lobule VI, dorsal caudal putamen (DCP), and dorsal rostral putamen (DRP), while our “cognitive” nodes are Crus I, Crus II, inferior and superior ventral striatum (VSi and VSs, respectively), dorsal caudate (DC), and ventral rostral putamen (VRP) as defined by prior work investigating connectivity (structural and functional) and functional activation patterns (Stoodley et al., 2012; Di Martino et al., 2008; Bernard et al., 2012; Salmi et al., 2010; Draganski et al., 2008). To do so, we averaged the beta values across their respective connections as a measure of connectivity strength to create indices of connectivity segregation for each participant. This resulted in average measures for three types of connections: motor-motor, cognitive-cognitive, and mixed (motor-cognitive). We then compared the degree of motor and cognitive segregation across both age groups using a 3 x 2 (connection type by age group) mixed model ANOVA. Follow-up within group repeated measures ANOVAs and paired samples t-tests were conducted in each age group separately.

## Results

### Cerebellar-Basal Ganglia Connectivity Patterns

First, we analyzed patterns of connectivity within each age group. In YA only, there were patterns of connectivity both within the cerebellar and basal ganglia seeds, but also between regions in a manner that is segregated based on functional topography (see Table 2; Figure 1a). Looking at connections between the cerebellum and basal ganglia, areas of the cerebellum involved in cognitive processing (Crus I and II) are correlated with regions in the basal ganglia (VSs and DC) that are functionally coupled with cognitive cortical regions (Di Martino et al., 2008) and are active during cognitive processing (Bernard et al., 2017). Similarly, areas associated with motor function in the cerebellum (Lobules V and VI) are correlated with areas predominantly connected to motor cortical regions in the putamen (VRP, DRP, DCP).

**Table 2:**
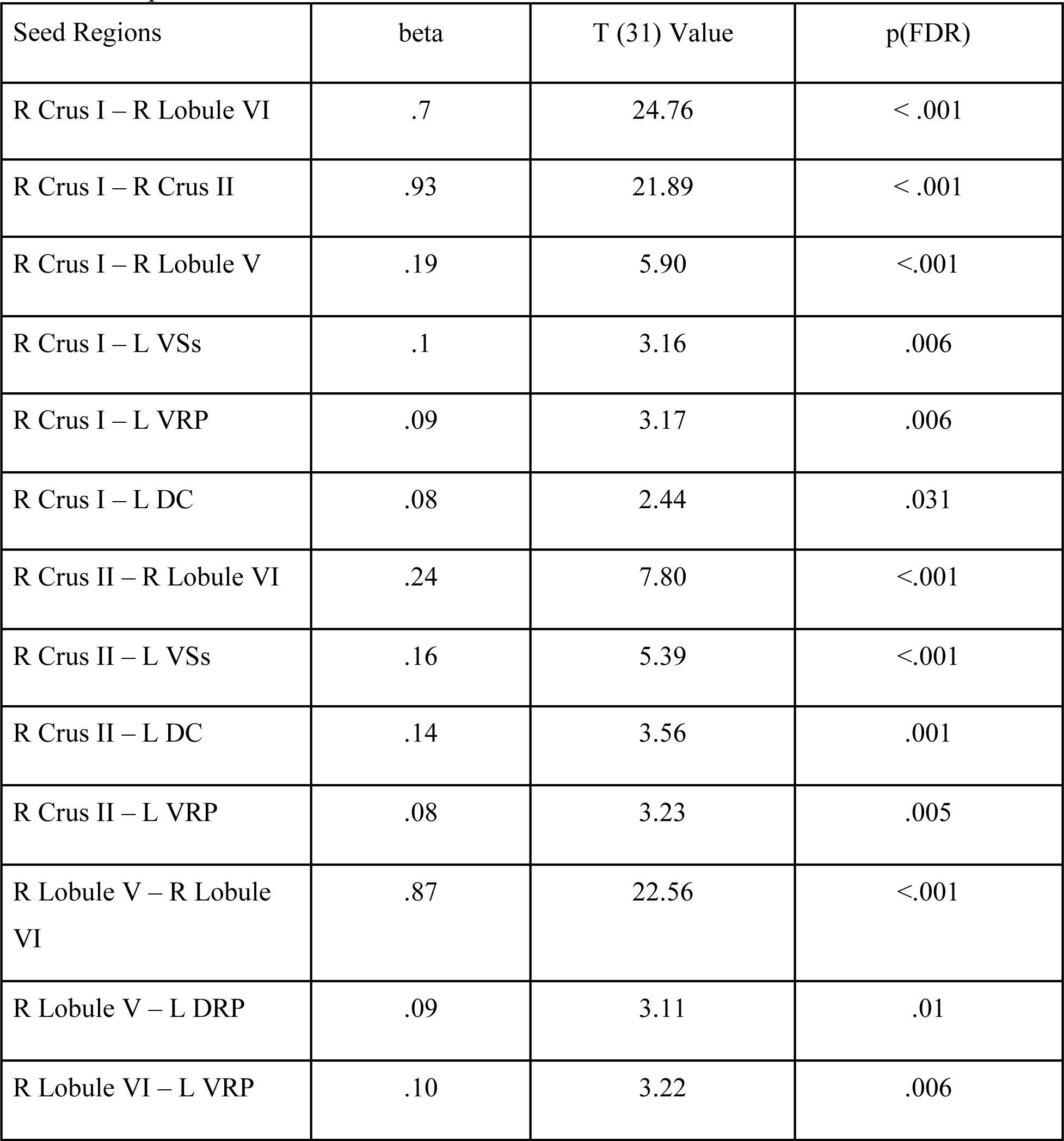
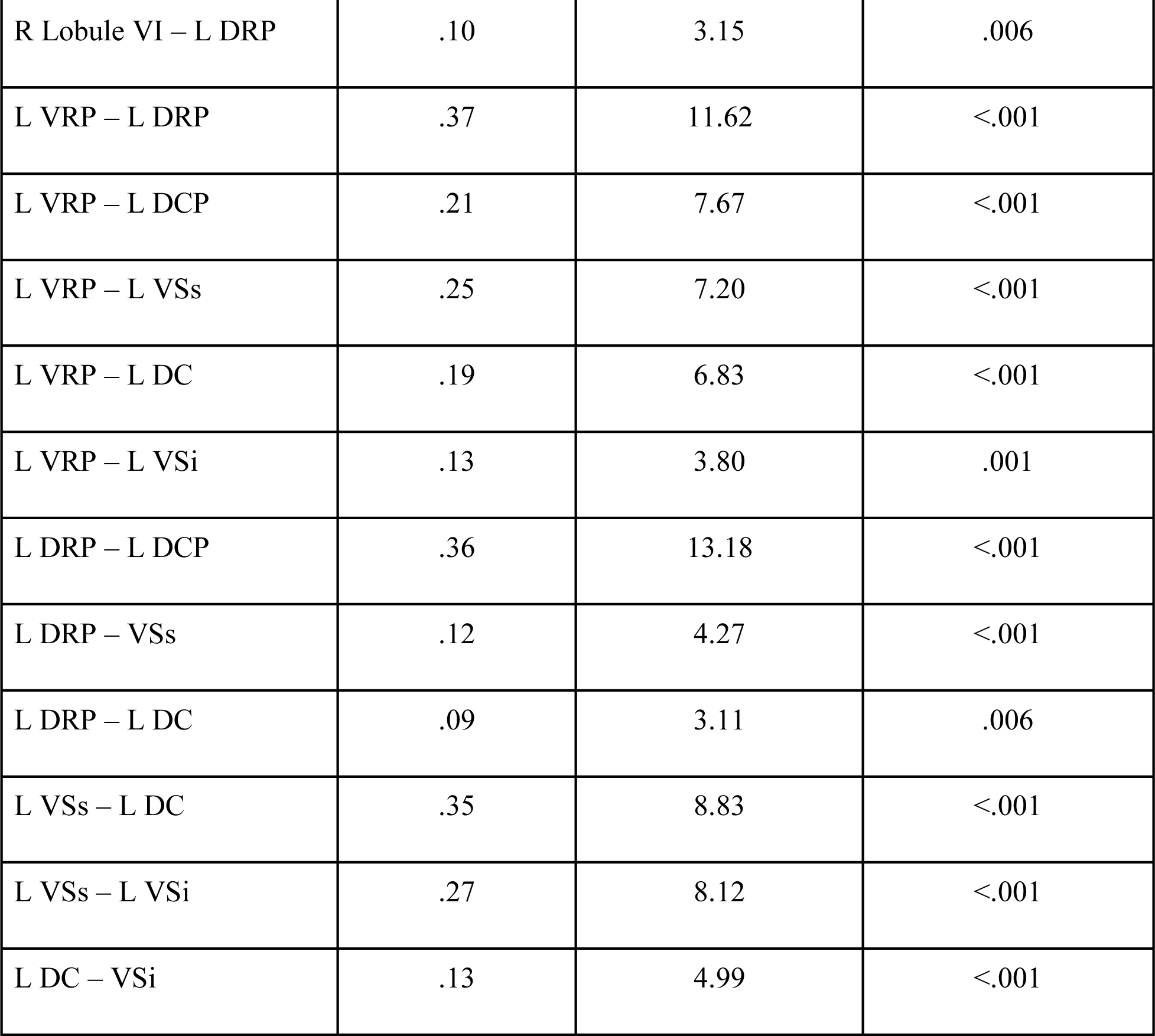
Significance tests of beta for ROI-ROI connections in YA only. Only significant ROI-ROI relationships are presented here. VSi: inferior ventral striatum; VSs: superior ventral striatum; DC: dorsal caudate; DCP: dorsal caudal putamen; DRP: dorsal rostral putamen; VRP: ventral rostral putamen.

| Seed Regions | beta | T (31) Value | p(FDR) |
| --- | --- | --- | --- |
| R Crus I – R Lobule VI | .7 | 24.76 | < .001 |
| R Crus I – R Crus II | .93 | 21.89 | < .001 |
| R Crus I – R Lobule V | .19 | 5.90 | < .001 |
| R Crus I – L VSs | .1 | 3.16 | .006 |
| R Crus I – L VRP | .09 | 3.17 | .006 |
| R Crus I – L DC | .08 | 2.44 | .031 |
| R Crus II – R Lobule VI | .24 | 7.80 | < .001 |
| R Crus II – L VSs | .16 | 5.39 | < .001 |
| R Crus II – L DC | .14 | 3.56 | .001 |
| R Crus II – L VRP | .08 | 3.23 | .005 |
| R Lobule V – R Lobule VI | .87 | 22.56 | < .001 |
| R Lobule V – L DRP | .09 | 3.11 | .01 |
| R Lobule VI – L VRP | .10 | 3.22 | .006 |
| R Lobule VI – L DRP | .10 | 3.15 | .006 |
| L VRP – L DRP | .37 | 11.62 | <.001 |
| L VRP – L DCP | .21 | 7.67 | <.001 |
| L VRP – L VSs | .25 | 7.20 | <.001 |
| L VRP – L DC | .19 | 6.83 | <.001 |
| L VRP – L VSi | .13 | 3.80 | .001 |
| L DRP – L DCP | .36 | 13.18 | <.001 |
| L DRP – VSs | .12 | 4.27 | <.001 |
| L DRP – L DC | .09 | 3.11 | .006 |
| L VSs – L DC | .35 | 8.83 | <.001 |
| L VSs – L VSi | .27 | 8.12 | <.001 |
| L DC – VSi | .13 | 4.99 | <.001 |

**Figure 1.**
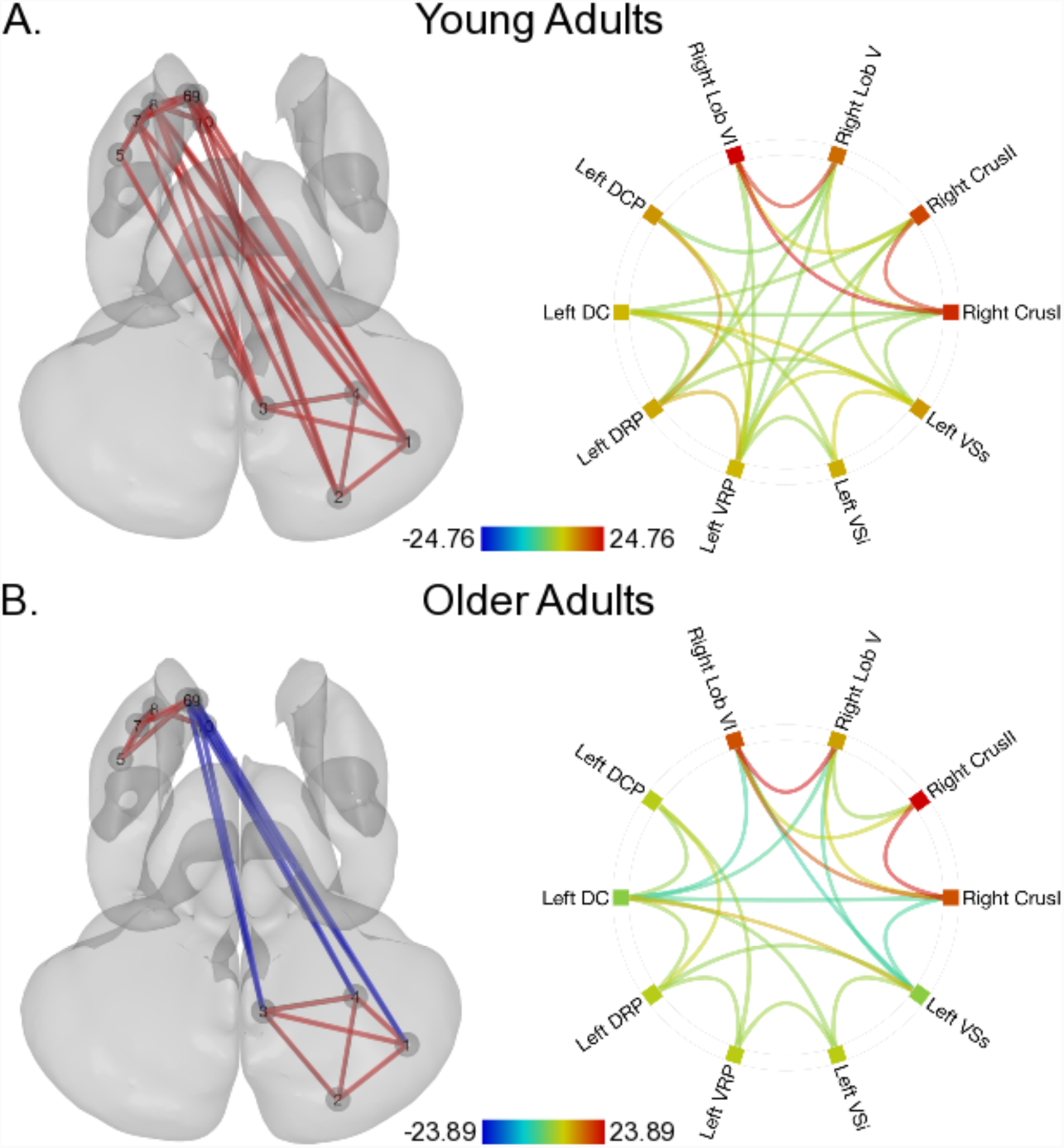
Significant ROI-ROI connections in **A.** young adults and **B.** in older adults. In both panels, the left image shows a three-dimensional extraction of the subcortical surfaces with the cerebellum at the bottom, and the basal ganglia at the top and more anterior. This image is presented such that the right hemisphere is on the right. The color of the connections is indicative of the direction of the associations (red: positive, blue: negative) though the color is absolute and provides no information regarding the degree of the associations. The connectome ring on the right provides an overview of the strength of the ROI-ROI connections within each age group. The color of the lines connecting regions represents the z-scores of the correlations between regions, and the color bar provides this range. The colored boxes in the ring are a combination of the effects across ROI-ROI connections and concatenate across the ROI-ROI effects. The warmer colors indicate more positive correlations between an ROI and the others in our network, whereas the cooler colors are suggestive of negative correlations. Numbers indicate the different cerebellar and basal ganglia seeds in the anatomical images. 1: Right CrusI; 2: Right CrusII; 3: Right Lobule V; 4: Right Lobule VI; 5: Left dorsal caudal putamen; 6: Left dorsal caudate; 7: Left dorsal rostral putamen; 8: Left ventral rostral putamen; 9: Left superior ventral striatum; 10: Left inferior ventral striatum.

### Age Differences in Cerebellar-Basal Ganglia Connectivity

A within-group analysis suggested that much like YA, OA show within cerebellum and within basal ganglia connectivity, as well as patterns of connectivity between subregions of these subcortical brain regions (Table 3; Figure 1b). Notably, while in YA we see positive correlations between the cerebellum and basal ganglia, the opposite is the case in OA. Patterns indicating significant *asynchrony* between these subcortical regions are present. For more information about the magnitude of the correlations between regions in YA and OA, beta values for all correlations are presented in Tables 2 and 3.

**Table 3:**
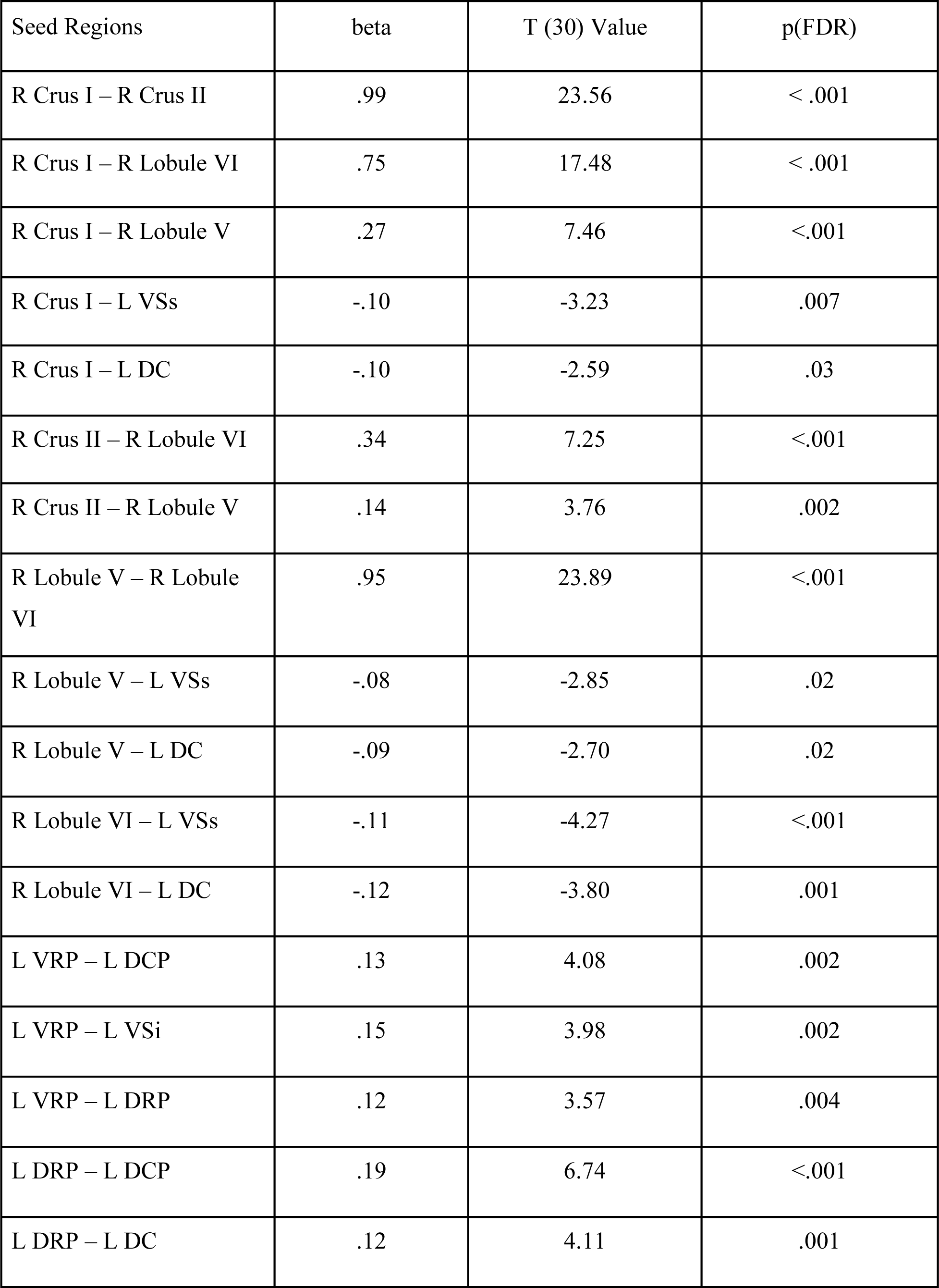
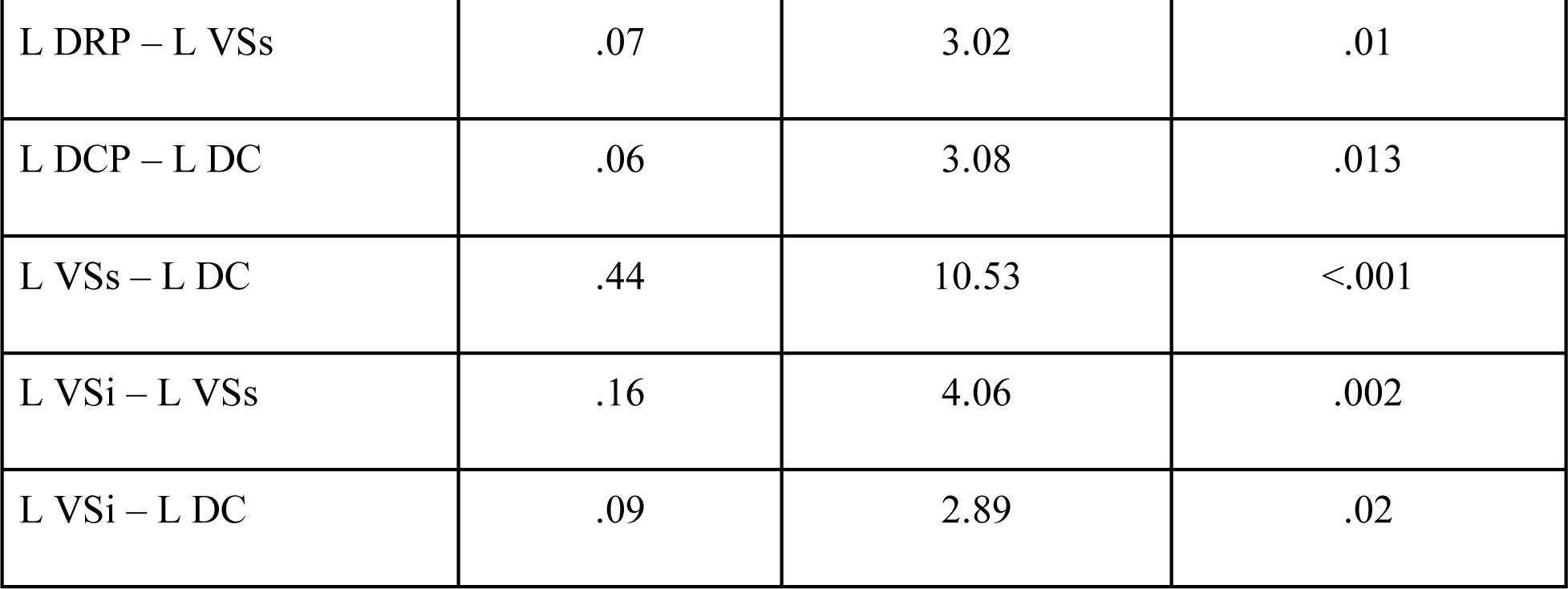
Significance tests of beta for ROI-ROI connections in OA only. Only significant ROI-ROI relationships are presented here. VSi: inferior ventral striatum; VSs: superior ventral striatum; DC: dorsal caudate; DCP: dorsal caudal putamen; DRP: dorsal rostral putamen; VRP: ventral rostral putamen.

Cerebellar-Basal Ganglia Connectivity
| Seed Regions | beta | T (30) Value | p(FDR) |
| --- | --- | --- | --- |
| R Crus I – R Crus II | .99 | 23.56 | < .001 |
| R Crus I – R Lobule VI | .75 | 17.48 | < .001 |
| R Crus I – R Lobule V | .27 | 7.46 | <.001 |
| R Crus I – L VSs | -.10 | -3.23 | .007 |
| R Crus I – L DC | -.10 | -2.59 | .03 |
| R Crus II – R Lobule VI | .34 | 7.25 | <.001 |
| R Crus II – R Lobule V | .14 | 3.76 | .002 |
| R Lobule V – R Lobule VI | .95 | 23.89 | <.001 |
| R Lobule V – L VSs | -.08 | -2.85 | .02 |
| R Lobule V – L DC | -.09 | -2.70 | .02 |
| R Lobule VI – L VSs | -.11 | -4.27 | <.001 |
| R Lobule VI – L DC | -.12 | -3.80 | .001 |
| L VRP – L DCP | .13 | 4.08 | .002 |
| L VRP – L VSi | .15 | 3.98 | .002 |
| L VRP – L DRP | .12 | 3.57 | .004 |
| L DRP – L DCP | .19 | 6.74 | <.001 |
| L DRP – L DC | .12 | 4.11 | .001 |
| L DRP – L VSs | .07 | 3.02 | .01 |
| L DCP – L DC | .06 | 3.08 | .013 |
| L VSs – L DC | .44 | 10.53 | <.001 |
| L VSi – L VSs | .16 | 4.06 | .002 |
| L VSi – L DC | .09 | 2.89 | .02 |

Also of interest are the differences as compared to YA. Our analyses demonstrated greater connectivity in YA relative to OA between the DRP and lobules V and VI. Lobules V and VI are strongly correlated with one another and implicated in motor networks (Kelly and Strick 2003; Krienen and Buckner 2009; Bernard et al. 2012). Given that the DRP is correlated with sensorimotor areas of the cortex (Di Martino et al., 2008), this demonstrates a decrease in communication between subcortical regions associated with motor processing networks. Indeed, while in YA we see strong correlations, in OA there are anticorrelations. There was also greater connectivity in YA relative to OA between the VSs and Crus I and Crus II, as well as between the DC and Crus I and Crus II. These regions of the basal ganglia and the cerebellum respectively are associated with cognitive networks and processing (Di Martino et al. 2008; Bernard et al. 2012; Stoodley et al. 2012; Bernard, Russell, et al. 2017). Interestingly, there is significantly higher connectivity between subregions within the basal ganglia in YA relative to OA; however, interactions within the cerebellum did not significantly differ between the two age groups. Overall, there is stronger connectivity in YA relative to OA within basal ganglia subregions as well as between basal ganglia and cerebellar areas involved in motor and cognitive functioning. This is likely driven in large part by the flip from synchrony to asynchrony in YA and OA. Table 4 provides the statistical information from this analysis, and the results are visualized in Figure 2. Notably, there was no greater connectivity in OA relative to YA between the cerebellum and basal ganglia. However, connectivity within the cerebellum between Crus II and Lobule V was significantly higher in the OA group relative to the YA group (seen in Table 4 as a negative beta value, given the design of our contrast).

**Table 4:**
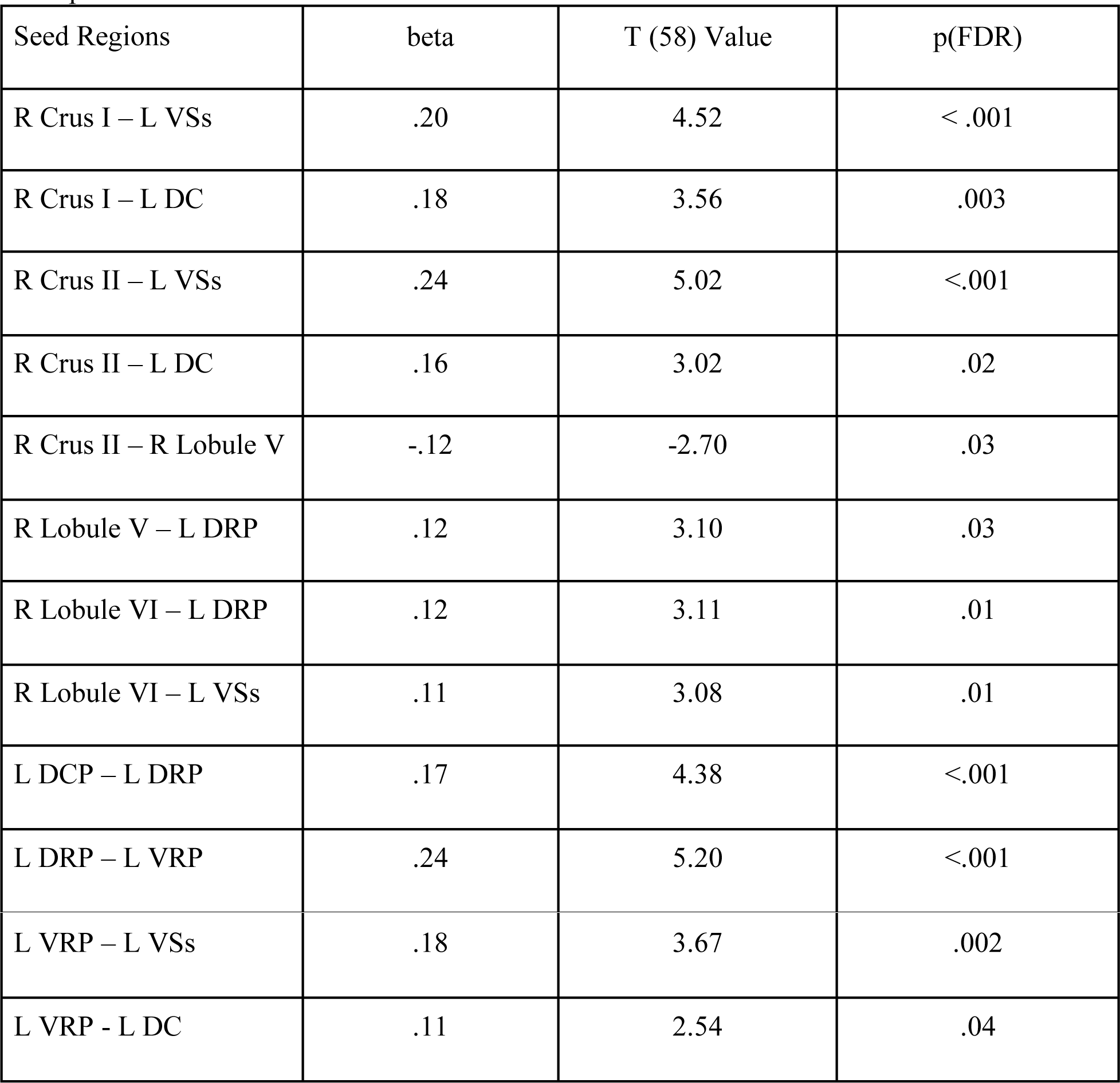
Significance tests of beta for regions in YA>OA contrasts. Only significant ROI-ROI relationships are presented here. VSi: inferior ventral striatum; VSs: superior ventral striatum; DC: dorsal caudate; DCP: dorsal caudal putamen; DRP: dorsal rostral putamen; VRP: ventral rostral putamen.

| Seed Regions | beta | T (58) Value | p(FDR) |
| --- | --- | --- | --- |
| R Crus I – L VSs | .20 | 4.52 | < .001 |
| R Crus I – L DC | .18 | 3.56 | .003 |
| R Crus II – L VSs | .24 | 5.02 | <.001 |
| R Crus II – L DC | .16 | 3.02 | .02 |
| R Crus II – R Lobule V | -.12 | -2.70 | .03 |
| R Lobule V – L DRP | .12 | 3.10 | .03 |
| R Lobule VI – L DRP | .12 | 3.11 | .01 |
| R Lobule VI – L VSs | .11 | 3.08 | .01 |
| L DCP – L DRP | .17 | 4.38 | <.001 |
| L DRP – L VRP | .24 | 5.20 | <.001 |
| L VRP – L VSs | .18 | 3.67 | .002 |
| L VRP - L DC | .11 | 2.54 | .04 |

**Figure 2.**
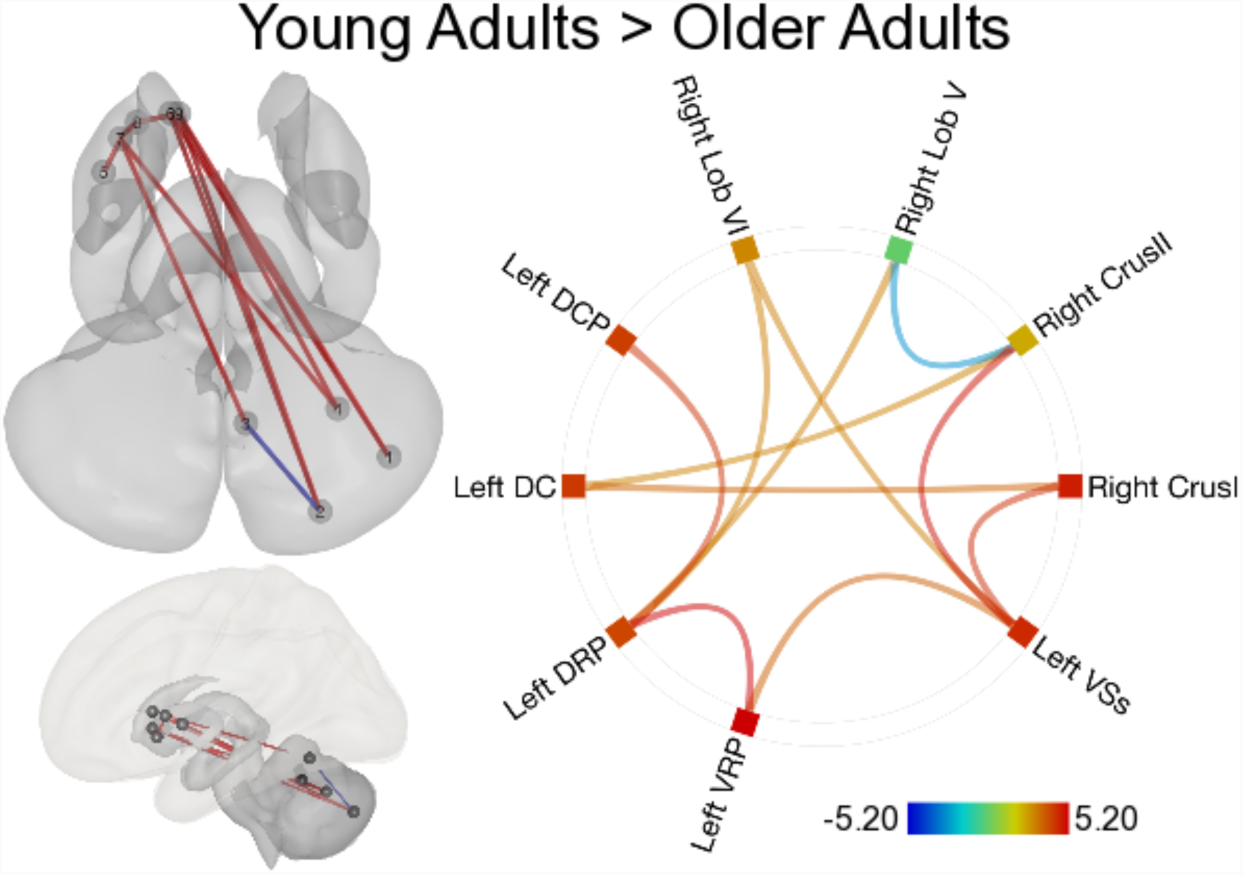
Age differences in cerebello-striatal connectivity. The top left provides an overview of the strength of the ROI-ROI connections demonstrating group differences in connectivity, paralleling the anatomical images from Figure 1. In all cases these data represent the contrast of young adults greater than older adults. The color of the connections is indicative of the direction of the associations (red: YA>OA, blue: OA>YA). On the bottom left the cerebellum and striatum shown in context with the cerebral cortex overlaid on top. In all cases these data represent the contrast of young adults greater than older adults. In the connectome ring, the color of the lines connecting regions represents the z-scores of the correlations between regions, and the color bar provides this range. The colored boxes in the ring are a combination of the effects across ROI-ROI connections and concatenate across the ROI-ROI effects. The warmer colors indicate more positive correlations between an ROI and the others in our network, whereas the cooler colors are suggestive of negative correlations. Numbers indicate the different cerebellar and basal ganglia seeds. 1: Right CrusI; 2: Right CrusII; 3: Right Lobule V; 4: Right Lobule VI; 5: Left dorsal caudal putamen; 6: Left dorsal caudate; 7: Left dorsal rostral putamen; 8: Left ventral rostral putamen; 9: Left superior ventral striatum; 10: Left inferior ventral striatum.

### Connectivity Strength and Behavior

Finally, to investigate the behavioral relevance of these subcortical functional interactions, we investigated the associations between striatal-cerebellar connectivity and cognitive function, as measured using the MOCA, and balance confidence as measured using the ABC Scale. One OA was excluded from the MOCA correlational analysis, having scored low on the MOCA, and while they were included in our connectivity analyses, they were not representative of the rest of our sample and had the potential to skew our findings. 1 OA and 3 YA were excluded from the ABC correlational analysis. The OA and 2 YA participants were excluded due to technical difficulties with the data, and 1 YA was not included due to improperly completing the ABC scale questionnaire, giving responses of 0, which would indicate that they are not confident at all in their balance. Not surprisingly, scores on the MOCA differed between YA and OA (YA= 27.56 ± 1.70; OA=25.87 ± 2.58; t(61)=3.08, p=.003). Due to the relatively limited range within each age group, we conducted these analyses with both age groups combined. Using a regression analysis implemented in CONN, we found that connectivity between Crus I and VSs was associated with MOCA score (*T*(61)=3.61, p_FDR_=.006, beta=0.04), suggesting that functional interactions between subcortical regions associated with cognitive cortical areas relate to general cognitive ability (Figure 3a). Because MOCA scores are highly correlated with age (*r*(63) = -.39, *p* = .002 in this sample), we repeated this analysis while controlling for age, and there were no significant correlations. Finally, we conducted the analyses in YA and OA separately. No association with MOCA scores were found in either group.

**Figure 3.**
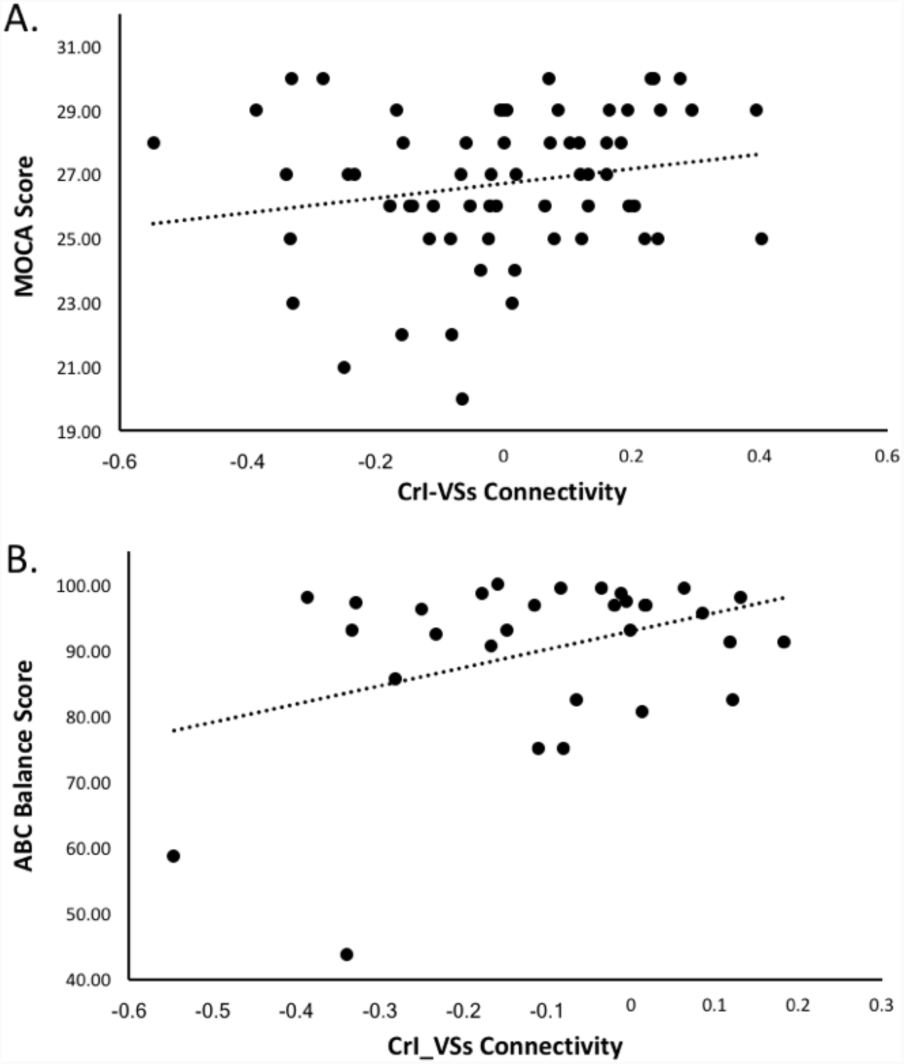
A. Scatterplot showing the relationship between MOCA scores and connectivity between Crus I and the superior ventral striatum (VSs). Data were collapsed across age groups. There were no significant relationships in YA and OA alone. **B.** Scatterplot showing the relationship between self-reported balance confidence as measured on the ABC Scale, and connectivity between Crus I and VSs in OA alone.

With respect to the ABC Scale, there were no significant age differences in self-reported balance confidence (YA=96.02 ± 5.74; OA=93.28 ± 12.83; t_(58)_=1.05, p=.296), consistent with our prior work using this scale in healthy older adults (Bernard & Seidler, 2013), though numerically balance confidence is rated lower in OA. In OA alone, connectivity between Crus I and VSs was significantly positively associated with self-reported balance confidence (t_(28)_=3.03, p_FDR_=.047, beta=.01; Figure 3b). This finding is consistent with our prior work looking at cerebellar network connectivity more broadly with respect to balance confidence in OA (Bernard et al. 2013). For both behaviors, these were the only significant associations to emerge, suggesting a relative degree of specificity in these results.

### Functional Circuit Segregation

The 3 x 2 (connection type by age group) ANOVA revealed several interesting findings (Figure 4). First, there was a significant main effect of circuit type (F(2,122)=4.022, p=.022, partial eta^2^=.062), and an interaction between circuit type and age group that approached the standard cut-off for significance (F(2,122)=2.438, p=.092, partial eta^2^=.038). Perhaps not surprisingly, there was a significant main effect of group as well (F(1,61)=20.66, p<.001, partial eta^2^=.253), driven by the shift from synchrony to asynchrony seen in YA and OA, respectively. A follow-up repeated measures ANOVA in the YA group alone further indicated a significant main effect of circuit type (F(2,62)=4.722, p=.012, partial eta^2^=.132), and follow-up paired t-tests demonstrated that there is no significant difference between the within-domain motor and cognitive circuits with respect to connectivity strength (t(31)=-.407, p=.687). Notably however, both the motor (t(31)=2.748, p=.01) and cognitive (t(31)=3.219, p=.003) circuits had stronger within domain connectivity when compared to the mixed cross-modal circuits, suggesting functionally determined circuit segregation in YA. Conversely, in OA, there was no significant main effect of circuit (F(2,60)=.934, p=.399, partial eta^2^=.03). Furthermore, for all paired t-tests comparing the circuit types to one another, there were no significant differences (for all analyses t(30)<1.32, p>.19). Critically, unlike in YA, we did not see the same degree of functional segregation in OA as connectivity strength for within modality functional circuits did not differ from the cross-modal mixed circuits.

**Figure 4.**
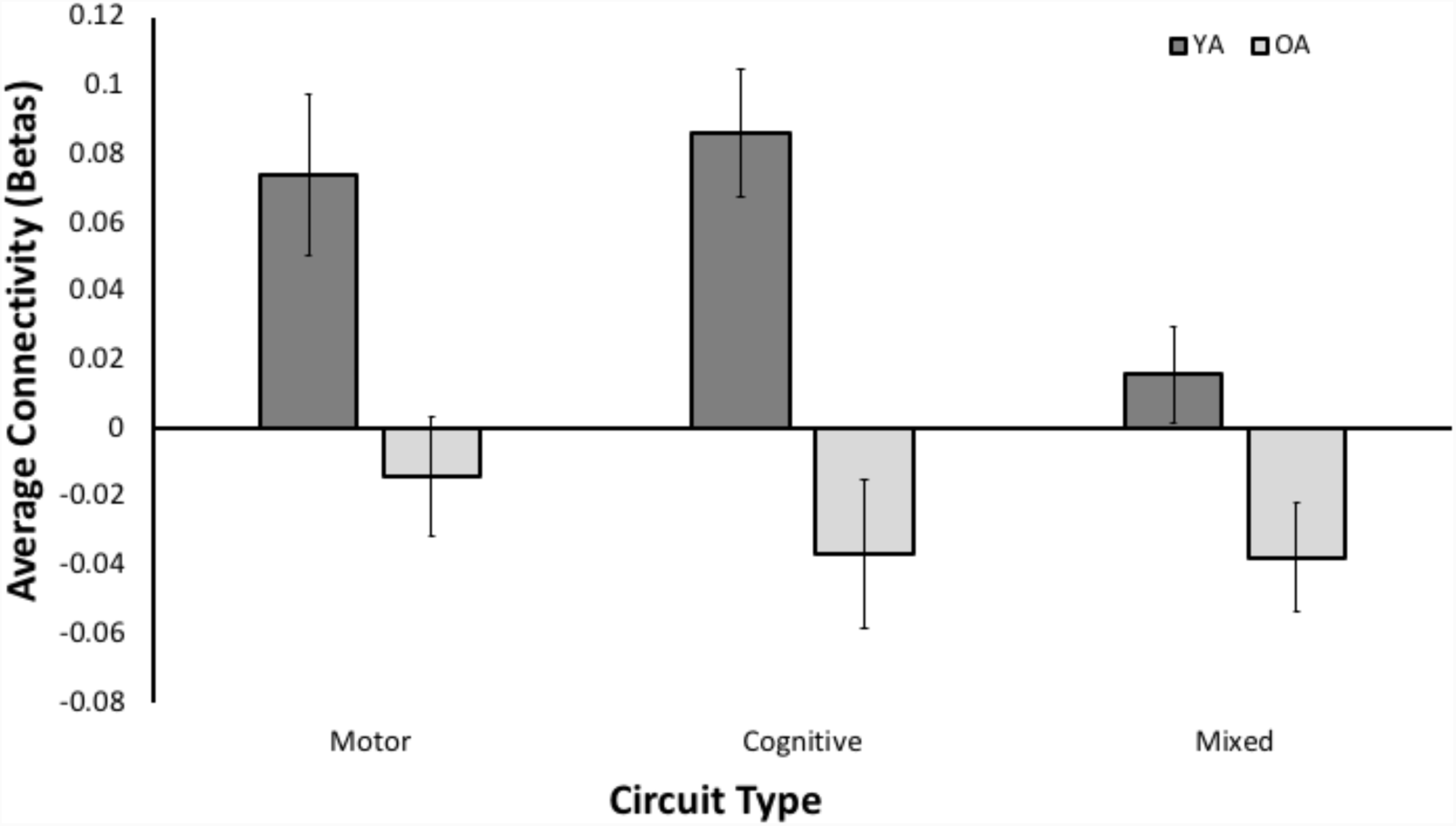
Age differences in functional segregation of cerebellar-basal ganglia circuits. In YA, both within domain circuits were more strongly connected (as determined by averaging beta values across connections) relative to mixed connections (motor-cognitive seed pairs). In older adults, in addition to the overall shift in connectivity to negative correlations, this pattern is no longer present. There are no differences between any of the different connection types.

## Discussion

Here, using an ROI-ROI based rs-fMRI analysis approach, we provide the first detailed mapping of striatal-cerebellar connectivity at rest in healthy humans supporting the notion of topographic organization of connectivity between these regions (Bostan and Strick 2018), and we have further demonstrated that there are age differences in these connectivity patterns. Broadly, connectivity in OA is weaker relative to that in YA, both within the striatum, and critically, between the cerebellum and striatum. Behaviorally, though data in this sample is limited, our results suggest that at least in terms of general cognitive function, there is an association with connectivity between the lateral posterior cerebellum (Crus I) and striatum (VSs). Taken together, these findings substantially broaden our understanding of the interactions between these two critical subcortical brain regions, both in healthy adulthood and in aging. Further, this is consistent with recent work suggesting that the basal ganglia and cerebellum form an integrated network with the cortex (Bostan and Strick 2018). This work has implications for our understanding of the role of striatal-cerebellar interactions from a processing perspective (i.e., what information is shared between these regions, how do they act together to produce behavior), in motor and cognitive declines in advanced age, and for age-related neurological disease, in particular Parkinson’s Disease.

### Cerebellar-Basal Ganglia Functional Connectivity Patterns

Work in non-human primates has demonstrated that the basal ganglia and cerebellum interact through disynaptic connections (Hoshi et al. 2005; Bostan and Strick 2010, 2018, Bostan et al. 2010, 2013). More recent work using diffusion tensor imaging has demonstrated parallel connections in the human brain as well (Pelzer et al. 2013; Milardi et al. 2016). While there have been suggestions in the broader rs-fMRI literature to indicate that there is functional coupling between the cerebellum and basal ganglia at rest (Di Martino et al. 2008; Bernard et al. 2012), a detailed investigation of the connections between individual lobules of the cerebellum and subregions of both the caudate and putamen had not been undertaken. Our results demonstrate that in healthy adults there is robust connectivity between ROIs in the cerebellum and basal ganglia and in patterns that are segregated based on similarity in functional connectivity patterns. For example, the dorsal caudate shows significant associations with both Crus I and II, but not lobules V and VI, while the dorsal and ventral rostral putamen seeds are strongly associated with Lobules V and VI, but not with Crus I and II. This parallels the relative functional segregation and topography of their cortical connectivity patterns with prefrontal and association cortices (dorsal caudate, Crus I, Crus II), and motor cortical regions (dorsal and ventral rostral putamen, Lobules V and VI) (Di Martino et al. 2008; Krienen and Buckner 2009; Bernard et al. 2012). This reciprocal communication between the regions indicates that the cerebellum and striatum are not just part of independent networks, but also exchange and integrate information with one another. Interestingly, in a recent comprehensive review of basal ganglia and cerebellar connectivity, Bostan and Strick (2018) suggest that the associations between these structures may underlie key cortical resting state networks. The segregated patterns of connectivity seen here in these pairwise analyses further support this notion.

### Age Differences in Cerebello-Striatal Connectivity

To this point, the primary focus with respect to the literature on the aging mind and brain has taken a cortical focus. While inroads have been made in our understanding of differences in subcortical network connectivity patterns (Bernard et al. 2013; Bo et al. 2014), the associations between the cerebellum and basal ganglia may be of particular importance. Both of these regions play a role in motor and cognitive behavioral performance (Stoodley and Schmahmann 2009; E et al. 2012; Stoodley et al. 2012; Bernard et al. 2013; Bernard, Russell, et al. 2017) and have distinct networks and connections with both prefrontal and association cortices and motor cortical regions (Kelly and Strick 2003; Di Martino et al. 2008; Draganski et al. 2008; Habas et al. 2009; Krienen and Buckner 2009; O’Reilly et al. 2010; Salmi et al. 2010; Bernard et al. 2012, 2014, 2016). Critically, both motor and cognitive performance are subject to age-related performance declines, and a more complete understanding of brain regions and their networks that contribute to these domains is necessary for understanding these changes. As such, both the cerebellum and basal ganglia are of great interest. In OA, there is evidence to suggest that there are differences in cerebellar volume and connectivity, both of which relate to behavioral performance (MacLullich et al. 2004; Raz et al. 2005b; Bernard and Seidler 2013; Bernard et al. 2013; Miller et al. 2013; Raz and Lindenberger 2013), and there are also known differences in basal ganglia functional connectivity across the lifespan with declines in advanced age (Bo et al. 2014). While our work has suggested that there are functionally relevant age differences in connectivity between the cerebellum and basal ganglia (Bernard et al. 2013), a detailed investigation of age differences in these connectivity patterns has not been undertaken to this point.

Here, we found that there are age differences in this circuit, broadly. Each cerebellar seed investigated showed significantly lower connectivity with striatal seeds. And while within region connectivity was not the primary focus of our investigation, there was markedly lower connectivity within the striatum in OA. Also of note is that within the OA alone, connectivity between the cerebellum and striatum was in the negative direction, opposite what was found in YA alone. Not only are the patterns of subcortical coupling significant different in YA and OA, they are divergent in terms of the direction of the associations. In YA there is strong coupling and synchrony between the cerebellum and basal ganglia, whereas in OA we see significant asynchrony suggesting an age-related shift in the interactions between these key subcortical regions. As noted above, Bostan & Strick (2018) have suggested that these interactions may be critical for broader resting state functional networks. Such a shift in the direction of the associations with age may have wide ranging impacts on functional connectivity more broadly in advanced age and may contribute to the age differences seen in the canonical cortical resting state networks (Andrews-Hanna et al. 2007; Wu et al. 2007; Damoiseaux et al. 2008; Onoda et al. 2012; He et al. 2014).

The shift in directionality with respect to cerebello-striatal connections is of great interest, particularly when considering underlying mechanisms and implications of these changes for our understanding of disease, particularly Parkinson’s disease. As suggested in our prior work which demonstrated age differences in cerebellar connectivity, including that with the basal ganglia which laid the foundation for our study here (Bernard et al. 2013), dopamine may be especially important. Even in healthy aging, there is a normative decrease in dopamine (McGeer et al. 1977; Fearnley and Lees 1991), and this change in dopamine may impact the functional interactions between these key subcortical regions. Further, in a study of healthy young adults, when administered l-dopa there is a significant increase in the connectivity between the cerebellum and basal ganglia, particularly in motor networks (Kelly et al. 2009). Notably, however, here we show this age-related asynchrony across connections. Thus, we speculate that during adulthood as dopamine naturally begins to decline, the nature of the interactions between the cerebellum and basal ganglia shifts from one of synchrony to that of asynchrony. When considering the implications of this finding for Parkinson’s disease, we suggest a “U-shaped” pattern of connectivity. That is, connectivity is high in healthy YA, lower and shifted in direction in healthy OA, and in Parkinson’s shifts to hyperconnectivity when off dopaminergic medication (e.g., Kwak et al. 2010; Festini et al. 2015). This hyperconnectivity shifts back more to normal patterns of connectivity when on medication (Kwak et al. 2010; Festini et al. 2015). An acknowledgement of this baseline type and degree of interaction in OA is important for our understanding of broader network connectivity, as well as the behavioral implications of these age differences.

In addition to our mapping of the resting state connectivity patterns between the cerebellum and striatum, we also investigated the behavioral relevance of these contributions. While our behavioral battery was limited, we demonstrated that connectivity between Crus I and the superior ventral striatum was positively associated with the MOCA. This parallels the functional specificity of the cerebello-striatal connections, as well as the functional topographies of these regions (Stoodley and Schmahmann 2009; E et al. 2012; Stoodley et al. 2012; Bernard, Russell, et al. 2017). Further, consistent with our prior work looking at cerebello-cortical connectivity in OA (Bernard et al. 2013), we found an association between self-reported balance confidence and connectivity again between Crus I and VSs. Our past work implicated Crus II and the caudate broadly (Bernard et al. 2013), and our replication here further underscores the importance of this circuit with respect to postural control. It is notable that these regions are associated more typically with cognition. However, there is a known increased reliance in cognitive function and circuits for postural control in OA (e.g., Verghese et al. 2002; Huxhold et al. 2006). Broadly, these findings provide support for the functional behavioral relevance of these connections, though future work with more extensive behavioral batteries is warranted. Further, it is important to note that the functional and computational roles of these regions in processing remains only theoretical to this point, with respect to the interactions between these structures (reviewed in Bostan & Strick 2018).

### Functional Circuit Segregation Differences with Age

In addition to investigating overall age differences in cerebellar-basal ganglia connectivity, we also investigated the relative segregation of purported motor and cognitive circuits between these two regions. Notably, we found that regions that shared similar patterns of cortical functional connectivity in a functionally segregated manner were more strongly connected than cross-modal connections. That is, regions of the cerebellum associated with motor function and motor cortical regions were more strongly connected to analogous basal ganglia regions, as compared to more cognitively associated regions. We found a parallel pattern for cognitively associated regions. Indeed, meta-analytic and connectivity research has demonstrated a relative degree of functional segregation and a functional topography within both of these subcortical structures (Stoodley and Schmahmann 2009; Bernard, Russell, et al. 2017). Consistent with this notion it is perhaps not entirely surprising to see an additional level of functional segregation with respect to connections between these subcortical regions. This also provides an additional mechanism for which cerebellar-basal ganglia connections may scaffold cortical networks.

Notably, this pattern of functional segregation was not present in OA. This marked lack of functional segregation suggests a degree of subcortical dedifferentiaion. At the cortical level, patterns of dedifferentiation, particularly in sensory and motor systems, have been observed in OA (Park et al. 2004, 2012; Carp et al. 2011; Bernard and Seidler 2012). That is, areas that are more functionally specialized in YA are less so in OA. Such dedifferentiation also impacts behavioral performance both within domain (Bernard et al. 2012), but also across domains (Park et al. 2010). For example, Park and colleagues used multi-voxel pattern analysis to investigate the specificity in visual processing for faces and houses in YA and OA. The degree of neural specificity was related to cognitive performance, wherein greater specificity was associated with poorer performance on multiple cognitive measures (Park et al. 2010). With that said, to our knowledge, this is the first demonstration of dedifferentiation at the subcortical network in OA. This decrease in network specificity and functional segregation in networks connecting these two key subcortical systems is of potentially great importance. While these segregated networks may provide key scaffolding for cortical networks in YA, the lack of segregation in OA may then negatively impact cortical functional connectivity, contributing perhaps at least in part, to the cortical functional connectivity differences seen in OA (e.g., Andrews-Hanna et al. 2007; Damoiseaux et al. 2008; Ferreira and Busatto 2013). Furthermore, though we did not include an extensive behavioral battery here, we speculate that this may also contribute, at least in part, to age differences in both motor and cognitive performance.

### Limitations and Future Directions

While our results greatly advance our understanding of the functional interactions between the cerebellum and basal ganglia in the human brain in YA and OA, there are also several limitations to this work. First, the behavioral testing battery was rather limited in our sample. While participants completed general questionnaire measures and the MOCA, they did not complete an extensive cognitive or motor task battery. As such, we are limited in our ability to investigate the behavioral implications of cerebellar-basal ganglia functional connectivity, and how this may relate to age differences in performance. While we demonstrated an association with MOCA scores in task-relevant regions, future work is needed to better understand the implications of these networks in both the motor and cognitive domain. Indeed, such work stands to be greatly illuminating for our understanding of behavioral differences in OA, as far less focus has been paid to subcortical regions and their networks.

In addition, we focused only on the caudate and putamen, using striatal sub-regions previously defined by Di Martino and colleagues (2008), though the animal work initially describing these connections implicated the globus pallidus *pars externa*, the subthalamic nucleus, and the cerebellar dentate (Hoshi et al. 2005; Bostan et al. 2010). However, we were concerned about possible challenges with the resolution of our rs-fMRI data, and we aimed for consistency in the size of our seeds. Seeds of different sizes will create an average time course across differing numbers of voxels, and as such, there is the possibility that this could impact our connectivity analyses. If the estimates of connectivity are created from substantially different numbers of voxels, as would be the case were we to use entire cerebellar lobules and small seeds in the basal ganglia, we may be unduly biasing and influencing our results. Further, there are known volumetric differences in the cerebellum lobules in older adults (Bernard and Seidler 2013). While we found that this did not impact connectivity in work investigating lobular cerebellar connectivity (Bernard et al. 2013), we do not know that this would necessarily be the case in this sample. Given the small size of the basal ganglia seeds, and the relatively large volume of the cerebellar lobules in their entirety, we matched the size of the seeds in this analysis. While it is possible to reliably measure connectivity from single-voxel seeds as we have done previously with the dentate nucleus (Bernard et al. 2014), we wanted to use seeds of the same size in our investigation here, and collect whole-brain data. Further, the 3.5mm radius employed here, modelled after Di Martino and colleagues (2008) would be too large to investigate the subthalamic nucleus or dentate without encompassing surrounding regions. Thus, we are limited with respect to our conclusions regarding functional interactions between the cerebellum and basal ganglia more broadly, as these smaller areas of the basal ganglia have not been included in our analyses. Future work with high resolution imaging, perhaps that which uses a limited number of high-resolution slices, is needed to look at these smaller regions more effectively.

On a similar note, we also limited out cerebellar seeds to only lobules V, VI, Crus I and Crus II. These regions show patterns of connectivity with primary motor, motor preparatory networks, and frontal-parietal cortical regions (e.g., Bernard et al. 2012; Krienen and Buckner 2009), many of which overlap with striatal-cortical resting state networks (Di Martino et al. 2008). While a comprehensive overview including all the lobules of the cerebellar hemispheres and vermis may be informative, it also creates issues with multiple comparisons (18 cerebellar regions, if only using one hemisphere and the vermis). We took a conservative statistical approach in our work here using an analysis-level correction at p_FDR_<.05, which took into account the number of pairwise connections in our model. While such a correction could be implemented in this analysis, we would be relatively underpowered given our sample size and the total number of seeds. As such, we focused on regions of the cerebellum that we predicted would show coherence with the striatal seeds used here, with the caveat that other cerebellar lobules may also show functional connectivity with the striatum and may be impacted by aging.

Finally, in our pre-processing and analysis, we used the Conn Toolbox, which includes normalization and processing as part of the pipeline. However, in some of our past work, we used a different approach including normalization with Advanced Normalization Tools (ANTS; Avants et al. 2008) in combination with in-house analysis scripts (Bernard and Seidler 2013; Bernard et al. 2013, 2014). This approach allowed for careful cerebellar normalization, and as past work has suggested, is more optimal for task-based functional analysis, as well as structural analysis (Diedrichsen 2006; Diedrichsen et al. 2009). Notably, in our more recent investigations we used CONN-based approach (Bernard et al. 2016; Bernard, Goen, et al. 2017), with results that consistently replicate work using ANTS. Critically, the first investigations of cerebello-cortical connectivity did not employ ANTS (Habas et al. 2009; Krienen and Buckner 2009; O’Reilly et al. 2010), and the robust patterns of connectivity have replicated across approaches. While there has been no direct comparison of cerebello-cortical connectivity using different normalization approaches, the pattern in the literature suggests that these networks are robust, and not as sensitive to differences across normalization approaches. Finally, it is important to note that here again we replicated and extended our prior work demonstrating age differences in cerebellar-basal ganglia connectivity (Bernard et al. 2013). This further supports the robust nature of these networks, and the ability to detect these patterns of connectivity across processing packages.

## Acknowledgments

The authors wish to thank all research participants for their time and effort taking part in this investigation. JAB was supported, in part, by the Brain and Behavior Research Foundation NARSAD Young Investigator Award as the Donald and Janet Boardman Family Investigator while this project was ongoing.

